# Neural signatures of stream segregation: From childhood to adulthood

**DOI:** 10.1101/2025.02.05.636482

**Authors:** Elena Benocci, Claude Alain, Axelle Calcus

## Abstract

When faced with noisy environments, listeners perform auditory scene analysis, which allows them to parse the auditory target from concurrent interferences. Stream segregation involves organizing similar sound waves into a coherent stream, while distinguishing dissimilar acoustic components and attributing them to distinct sources. Two event-related potential components have been identified as “neural signatures” of stream segregation: the Object-Related Negativity (ORN) and the P400. Our study aims to examine (i) the maturation of neural and behavioural correlates of stream segregation and (ii) the development of the relationship between stream segregation and speech perception in noise. ORN/P400 were recorded while 8-23 year-olds performed an active stream segregation task based on temporal coherence. Participants also performed speech identification in noise tasks (behaviourally). Behavioral results indicate an improvement in both stream segregation and speech perception in noise from childhood to adulthood. Amplitude of the ORN (but not P400) decreases, and latency of both ORN and P400 decreases throughout development. Critically, P400 amplitude significantly predicts stream segregation performance. Overall, our results suggest that the neural mechanisms underlying stream segregation follow a prolonged maturation trajectory, and support the progressive maturation of auditory scene analysis and speech perception in noise.

**RESEARCH HIGHLIGHTS:**

- Neurophysiological indices of stream segregation mature from childhood to adulthood
- Neural indices of top-down attentive processing of complex auditory sequences predict stream segregation, irrespective of the listeners age
- In adults (but not in children or adolescents), stream segregation predicts speech perception in noise
- Stream segregation may thus operate as a bottleneck to speech in noise difficulties in children and adolescents

## INTRODUCTION

From playgrounds to classrooms and to sports practice, children and adolescents routinely navigate complex auditory environments. In everyday environments, sound waves from various sources are mixed before reaching the ears. Listeners must disentangle individual sounds from the mixture, performing the auditory scene analysis (ASA; Bregman, 1990, 2015). Adequate analysis of complex auditory scenes relies upon the listeners’ ability to parse acoustic events into different streams (Moore & Gockel, 2002). Newborns perform auditory stream segregation from birth, albeit less efficiently than adults (Folland et al., 2012; McAdams & Bertoncini, 1997). How auditory streaming develops in the first decades of life remains poorly understood.

### Development of stream segregation

Canonical studies of stream segregation use sequences of tones organized temporally in repeated A-B-A—A-B-A patterns, where A and B represent successive tones of different frequencies and – represents a silence (Miller & Heise, 1950; Van Noorden, 1975). Adult listeners are asked to report whether they heard one or two streams as a function of parametrical changes of the sequences (e.g., A and B frequency distance, repetition rate, etc). When listeners report hearing two streams, they are effectively experiencing stream segregation: they parse the sequential auditory events into distinct streams. At a given presentation rate, increasing the frequency distance between A and B typically increases the likelihood that listeners will report hearing distinct sound streams. Early developmental studies suggest that the frequency distance needed for children to experience sequential stream segregation is much larger than adults’ (Sussman et al., 2007; Sussman & Steinschneider, 2009). In fact, it keeps decreasing from childhood to adolescence, until 15 years at least (Sussman et al., 2015).

However informative, this canonical paradigm is blind to an important aspect of auditory scene analysis, in which auditory objects tend to occur simultaneously (for a review, see Micheyl & Oxenham, 2010). Another paradigm was thus developed to evaluate segregation of concurrent sounds. Listeners were presented with harmonic complex sounds, of which one tonal element was mistuned (Moore, Glasberg and Peters, 1986) or delayed (Hedrick & Madix, 2009) – manipulations that contribute to segregation into distinct auditory objects. The larger the mistuning/the delay, the more likely adults will report hearing two auditory objects, hence experiencing concurrent stream segregation. Again, children require larger changes in frequency to detect a mistuned harmonic than adults (Alain et al., 2003). How concurrent stream segregation develops at adolescence remains unknown.

Specific neurophysiological markers of stream segregation have been associated with each paradigm. First, the mismatch negativity (MMN) has been used as an index of sequential segregation. The brain generates a MMN when it processes a difference between an unexpected auditory stimulus (the deviant) and the neural representation of an expected (standard) stimulus (for a review, see Näätänen et al., 2012). In a clever adaptation of this paradigm, adults were presented with sequences of interleaved sounds of different frequencies. An infrequent (louder) deviant was placed in the sequence, but it would only be processed as such when the frequency difference between successive sounds was large enough (hence eliciting the perception of two streams). In sequences processed as containing two auditory streams, an MMN would thus appear in response to the infrequent tone (Sussman, Ritter and Vaughan, 1999). Similar results were reported in newborns, revealing an innate ability to parse auditory scenes (Winkler et al., 2003). Children (7-10 years) also showed an MMN in response to infrequent tones placed in sequences that promote stream segregation (Sussman et al., 2001; Lepistö et al., 2009). The MMN amplitude is thought to be similar in children (9-12 years) and adults if they are actively engaged in detecting the infrequent target, but not if they passively listen to the sequences (Sussman & Steinschneider, 2009).

The object-related negativity (ORN) and P400 are event-related potentials (ERPs) that index listeners’ processing of two concurrent auditory objects (Alain, Arnott and Picton, 2001; for a review, see Gohari et al., 2022). The ORN is an early, pre-attentive, negative response with a fronto-central distribution (Alain & Izenberg, 2003; Alain & McDonald, 2007; McDonald & Alain, 2005). When participants are actively engaged in a segregation task, the ORN is followed by a positive component (P400) that displays a centro-parietal distribution (Alain et al., 2001; Alain, Schuler and McDonald, 2002). Both ORN and P400 are typically elicited by comparing the neurophysiological response evoked by an in-tune complex harmonic to that of a harmonic complex that contains a mistuned component. When presented with such a mistuned component, newborns and infants show an ORN, again supporting the hypothesis of an innate ability to process concurrent streams (Bendixen et al., 2015; Folland et al., 2015; Mehrkian et al., 2022). Surprisingly, presented with the same stimuli, 8-13 year-old children displayed a *larger* ORN than adults (Alain et al., 2003) – but no significant P400 (Alain et al., 2003; Mehrkian et al., 2022). Together, neurophysiological studies thus suggest that children (∼8-12 years) show immature neurophysiological responses in situations of either sequential or concurrent streaming (Sussman et al. 2001; Alain et al., 2003). How these responses develop at adolescence has remained so far unexplored.

A major limitation of these early studies is their focus on *either* sequential or concurrent stimuli. In everyday communication, listeners seek to understand a given speaker speech stream in the presence of other, interfering speakers. Concurrent speech signals are broadband, often highly correlated, and overlapping with one another. In such situations, temporal coherence between different acoustic cues of the auditory objects is crucial for auditory segregation (Elhilali et al., 2009). The “stochastic figure-ground paradigm” (SFG) was developed to evaluate streaming abilities based on temporal coherence, hence requiring listeners to perform both sequential *and* concurrent streaming (Teki et al.,, 2011; Teki et al., 2013). In this paradigm, listeners have to detect a temporally coherent figure presented against a background of randomly varying tones. Adults are remarkably sensitive to the appearance of such figures in stochastic noise backgrounds: discrimination performance even improved as figure coherence/duration increased (Teki et al., 2013). Both ORN and P400 have been recorded in adults as they performed the SFG task (Tóth et al., 2016). Some studies found significant relationships between ORN/P400 amplitude and behavioural streaming performance (Alain et al., 2001, 2002; Zendel & Alain, 2013; Tóth et al., 2016), but not all (Lipp et al.,, 2010; Lodhia et al., 2014). So far, the causal relationship between ORN/P400 amplitude and streaming performance remains debated.

To sum up, both behavioural and neurophysiological studies indicate that stream segregation remains immature until adolescence. Yet the exact age of maturation and the neural processes underlying maturation remains unknown. Therefore, the main aim of this study is to investigate the development of both behavioural and neurophysiological indices of stream segregation from childhood to adulthood. A second objective of this study is to investigate the link between neurophysiological indices and performance in a stream segregation task that relies upon temporal coherence.

### Development of speech perception in noise

In everyday life, listeners are not presented with pure or complex tones, but rather with speech signals uttered in the presence of environmental noise or competing speech. Whereas the former is thought to interfere with the speech signal at the peripheral level (*energetic masking*), the later typically affects auditory processing at the central level (*informational masking*; for a review, see Durlach, Mason, Kidd and Arbogast, 2003). In adults, speech intelligibility is worse in a background of interfering speakers than in (spectrally identical) speech-shaped noise (SSN) (Arbogast, Mason and Kidd, 2002; Brungart et al., 2001; Brungart, 2001; Kidd, Mason and Gallun, 2005), likely due to the added cognitive contribution inherent to speech stimuli (Mattys et al., 2012). Note that in the presence of only one interfering speaker, syllable identification tends to be better than in the presence of a spectrally identical SSN (Wang & Xu, 2024). This benefit likely stems from the possibility to glimpse in brief dips of a 1-talker masker to improve the identification of the speech target (Simpson & Cooke, 2005).

In children, the peripheral auditory system is functionally mature at birth (Werner, 2017), but complex auditory processes such as speech-in-noise (SiN) perception follow a protracted developmental trajectory (for a review, see Leibold & Buss, 2019). The exact timing of maturation for SiN perception remains uncertain, but appears to be dependent upon the nature of the interference induced by the background noise. In the presence of SSN, speech intelligibility seems to be mature between 7 and 11 years of age (Cabrera et al., 2019; Corbin et al., 2016; Evans & Rosen, 2021; Koopman, Houtgast and Derschler, 2008) - although adolescents still improve by about 0.2 dB SNR per year between 12 and 17 years of age (Jacobi et al., 2017). In the presence of one or two interfering speakers, speech intelligibility gradually improves until 13 (Buss et al., 2017; Baker, Buss et al., 2014; Corbin et al., 2016; Leibold & Buss, 2013) or even 16 years of age (Buss et al., 2017; Calandruccio et al., 2020; Wightman & Kistler, 2005). In the presence of multiple speakers (8- to 20-talker babble), speech intelligibility improves from childhood to late adolescence (Bonino, Leibold and Buss, 2013; Elliott, 1979; Ginzburg et al., 2024; Lewandowska et al., 2021). In short, speech intelligibility in the context of energetic masking appears to reach maturity by late childhood. Intelligibility in backgrounds that induce mostly informational masking appear to follow a protracted developmental trajectory, from childhood to (late) adolescence.

### Relationship between streaming and SiN perception during development

Very few studies have investigated the relationship between streaming and SiN perception throughout development. In adults, behavioural and neuroimaging studies concur to support a relationship between stream segregation and SiN perception (Holmes & Griffiths, 2019; Holmes et al., 2021). Our recent study examined the direction of this relationship throughout development (Benocci & Calcus, 2024). Children, adolescents and adults were presented with both the SFG task and speech perception in noise tasks (in the presence of SSN and one interfering talker). Children had more difficulties than adolescents on the SFG task (adolescents’ performance did not significantly differ from adults’). The same developmental improvement was observed in a consonant identification task in the presence of SSN (children being poorer than adolescents, who did not differ from adults). In the presence of one interfering talker, we observed a gradual improvement until adulthood. Using structural equation modelling, stream segregation predicted SiN perception (irrespective of the noise condition). Age significantly contributed to both stream segregation and SiN perception. Remarkably, musical abilities significantly contributed to SiN perception through a mediating effect of stream segregation.

This finding was in line with the bulk of the literature indicating a musician advantage for SiN perception in adults (Besson, Chobert and Marie, 2011; Parbery-Clark, Skoe and Kraus, 2009; Parbery-Clark et al., 2011; Parbery-Clark et al., 2012; Slater & Kraus, 2016; Kalplan et al., 2021; see Hennessy, Mack and Habibi, 2022 for a meta-analysis) – although some studies did not find such a musician advantage (Boebinger et al., 2015; Coffey et al., 2017; Madsen, Whiteford & Oxenham, 2017; Ruggles, Freyman and Oxenham, 2014; Yeend et al., 2017). Similarly, adult musicians seem to exhibit better stream segregation abilities than nonmusicians (Alain et al., 2014; Zendel & Alain, 2009; Zendel & Alain, 2014). In adolescents, music lessons in high-school are thought to support neural processing of speech in quiet (Tierney, Krizman and Kraus, 2015). Two years of active musical training in elementary school even led to significant benefits for SiN perception (Slater et al., 2015). Note that all the studies mentioned here compared musicians to non-musicians, using a dichotomous categorization that may not reflect the full range of musical abilities within the population (Law & Zentner, 2012). Such a categorical approach is particularly problematic in developmental studies, where the duration of musical training (which is typically used as a criterion to distinguish between musicians and non-musicians) is necessary limited in the younger participants. Therefore, measuring musical abilities on a continuous scale might offer a more accurate view of the relationship between music and speech processing (Zentner & Strauss, 2017).

### The present study

The objective of this study was twofold: investigate (i) the development of, and (ii) relationship between behavioural and neurophysiological indices related to stream segregation and SiN perception, from childhood to adulthood. Regarding the first objective, we predict a significant decrease in ORN amplitude from childhood to adulthood. The presence/morphology of a P400 in childhood/adolescence remains debated, and will be explored here. Regarding the second objective, adult studies show contradictory results regarding the relationship between ORN/P400 and streaming performance. To the best of our knowledge, no study has investigated the relationship between ORN/P400 and SiN perception. Therefore, we explore these relationships throughout development. A third, exploratory, hypothesis is that musical abilities could influence behavioural and neurophysiological indices related to stream segregation and SiN perception.

## METHOD

### Participants

Seventy-eight French natives were recruited for this study, including 24 children (aged 8-12 years, 12 males), 27 adolescents (aged 13-17 years, 14 males), and 27 adults (aged 18-22 years, 11 males). EEG data from 10 participants were excluded due to noisy recordings. The final sample thus included 23 children, 22 adolescents, and 25 adults. All participants had normal audiometric thresholds (≤ 20 dB HL) as measured at octave intervals from 0.25 to 8 kHz. Cognitive abilities were evaluated using the WISC-V (children/adolescents) or the WAIS-IV (adults). Specifically, working memory was assessed using the Working Memory Index and fluid reasoning was assessed using the Matrix Reasoning. Two adolescents who did not exhibit normal working memory and reasoning abilities for their age were excluded. The normalized cognitive results by age group for the final sample is available in Table 1. All included participants were non-musicians: they had less than three years of formal musical training (Parbery-Clark et al., 2011; Slater & Kraus, 2016; Yoo & Bidelman, 2019). None of them reported any history of neurological, hearing, or developmental disorders. Two participants were left-handed, as confirmed by the Edinburgh Handedness Inventory (Oldfield, 1971). Prior to the study, all participants (and their legal guardians, when needed) provided written consent. The protocol of the study was approved by the ethical committee of the Hôpital Universitaire des Enfants Reine Fabiola (CEH n°48/21).

### Behavioural testing

#### Musical abilities – MICRO-PROMS

The MICRO-PROMS battery is an objective tool designed to evaluate musical perceptual abilities (Strauss et al., 2023). This streamlined version of the original PROMS battery (Law & Zentner, 2012) uses a three-interval forced-choice procedure to assess discrimination abilities across various musical dimensions: accent, melody, pitch, rhythm, tempo, tuning, and timbre. In each trial, participants heard two repetitions of a reference musical stimulus, followed by the target stimulus. Participants had to judge whether the reference stimulus and the target were similar or different. The target stimulus could only differ from the reference stimulus along one of the musical dimension at a time. In total, 18 trials were presented to the participants. This task was presented through Sennheiser (HD300) headphones at an intensity of 73 dB SPL.

#### Speech perception in noise

The stimuli were the same as in Benocci & Calcus (2024). Participants were presented with 32 vowel-consonant-vowel (VCV) logatomes, which comprised two repetitions of 16 possible/aCa/ logatomes, spoken by a French female voice. The consonant was one of the following: /p, t, k, b, d, g, f, s, ∫, m, n, r, l, v, z, j/. Logatomes were presented in silence and in two different noisy backgrounds: an interfering speaker (1-talker) and SSN. Although a single interfering talker is not the most disruptive for speech perception (Simpson & Cooke, 2005), it was selected because it is somewhat comparable to the stochastic figure-ground task of discriminating a coherent target figure (here: a logatome) from a similar-sounding background.

This masker was composed of a male speaker reading French press articles. Pauses longer than one second, pronunciation errors, and proper nouns were edited out (without being noticeable by the listeners). SSN maskers were then generated from the 1-Talker condition using a fast Fourier transform. This process examined the power and phase spectrum of the original recording. A new signal was created with the same power spectrum but randomized phases, resulting in the SSN.

Each target logatome was approximately 500 milliseconds long and both types of background noise lasted two seconds. The logatomes started randomly between 200 milliseconds and one second after the start of the noise. All audio files were standardized to the same root-mean-square level. In noisy conditions, the signal-to-noise ratio (SNR) was set at -5 dB. The target logatomes and the noise were played simultaneously to both ears. This task was implemented via the Gorilla online testing platform (Anwyl-Irvine et al., 2020). Sounds were presented through Sennheiser (HD300) at an intensity of 73 dB SPL.

### Neurophysiological testing: *Auditory stream segregation*

Stream segregation was evaluated using the stochastic figure-ground task as described by Teki et al. (2013), with parameters that led to the highest discrimination scores in Experiment 1. Trials consisted in 264 2-s long sequences of 5 to 15 50-ms simultaneous pure tones with random frequencies ranging from 179 to 7246 Hz (log-distribution), forming a background of inharmonic chords, with no inter-chord interval. In 2/3 sequences, a subset of eight pure tones were repeated over seven chords, constituting a 350-ms figure (SNR = 0 dB). The figure started between 750 and 1000 ms after the sequence onset. A third of the sequences did not contain any figure, consisting merely in the stochastic background (henceforth: control sequences). All sequences were presented binaurally at 75 dB SPL (ER-2, Etymotic research).

Participants were asked to indicate whether they heard a figure in each trial. They were first given examples of the figure alone and background, followed by ten practice trials. Feedback was provided throughout the entire task. Performance was measured by calculating hit rates (correct detections of the figure when present) and false alarm rates (incorrect reports of a figure when only the background was present). These rates were then used to compute the d’ sensitivity index, determined by the difference between the z-scores of the hit rates and the false alarm rates, as described by Macmillan and Creelman (2005).

The neuroelectric brain activity was continuously recorded during this task using a BioSemi Active Two system at a sampling rate of 2048 Hz, from 64 scalp locations, according to the international 10–20 system (Jasper, 1958). Additional electrodes were placed on each mastoid and recordings were re-referenced offline to the average of activity at the mastoid electrodes. Electrode offsets were consistently < 50 mV. Preprocessing was carried out with MATLAB 2021b (MathWorks) using custom scripts based on EEGLAB (Delorme & Makeig, 2004). The recorded data was band-pass filtered between 1 and 30 Hz (zero-phase, finite impulse response, -6 dB/octave). Event-related potentials were obtained by creating 1000-ms long epochs that started 50 ms before figure onset. In control sequences, epochs started 700 ms after sequence onset. All 264 epochs were baseline-corrected using the mean value from -50 to 0 ms. Epochs exceeding ± 100 μV were rejected.

Difference waveforms were obtained by subtracting the ERPs corresponding to control sequences to those of figure sequences, averaged across a subset of electrode. The subset of electrodes were defined by visually inspecting the voltage maps of the responses (see Figure 1 for group averages). For the ORN, we selected electrodes F1, F2, F3, F4, FC1, FC2, FC3, FC4, Fz, and FCz. For the P400, we selected electrodes CP1, CP2, P1, P2, Pz, CPz, and POz.

**Figure 1.**
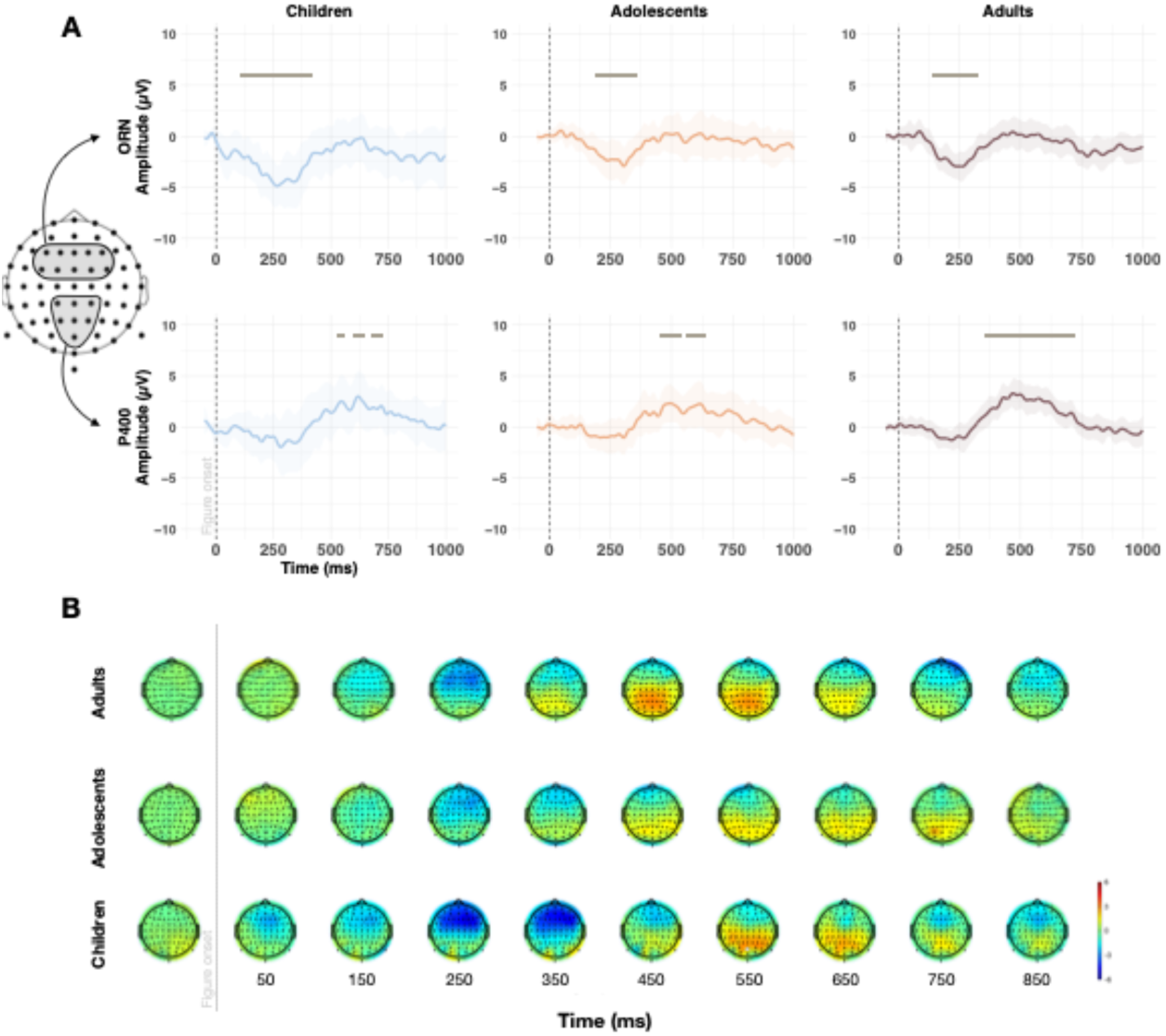
A: Grand averages of the ORN (top, from fronto-central electrode cluster) and P400 (bottom, from parietal electrode cluster) by age group, including all participants. Periods of a significant (*p* < 0.01 for > 20 ms) ORN and P400 are shown by a horizontal line, as a function of age group. B: Voltage maps show the mean differential response (figure trials – control trials) during the 0-850 ms post-stimulus time window.

The ORN and P400 were statistically assessed in three ways. First, we tested whether ORN/P400 were present at the group level. To do so, we conducted point-to-point comparison of each group differential wave amplitude to determine the latency period over which the waveforms were significantly smaller (ORN) or larger (P400) than zero, if any. One-sided *t*-tests were computed between 70 and 500 ms (ORN) and between 300 and 800 ms (P400) post figure onset. Because adjacent points in the waveform are highly correlated, potentially leading to spurious significant values in short intervals, an ORN/P400 was considered present when p < 0.01 (one-tailed) over a period of ≥ 20 ms at adjacent time-points (Kraus et al., 1993; McGee et al., 1997). Second, we aimed at computing the individual amplitude and latencies of ORN/P400 in each participant. Individual latencies corresponded to the timing of the minimum/maximum amplitude within the ORN/P400 (respectively) significant time window observed in the corresponding group. Individual amplitudes were computed as the average amplitude recorded in an 80 ms window around the individual peak. Finally, we sought to determine how many individuals showed significant ORN/P400 within each group. Indeed, differences in amplitude/latency could merely be due to the proportion of individuals showing each specific component within each group. Therefore, one-sided *t*-tests (p < 0.01 for ≥ 20 ms at adjacent time-points) were performed again, on *individual* differential waves (averaging responses from the same electrodes as those used to look at the grand average ORN/P400), details by age group are available in Table 2.

**Table 2:**
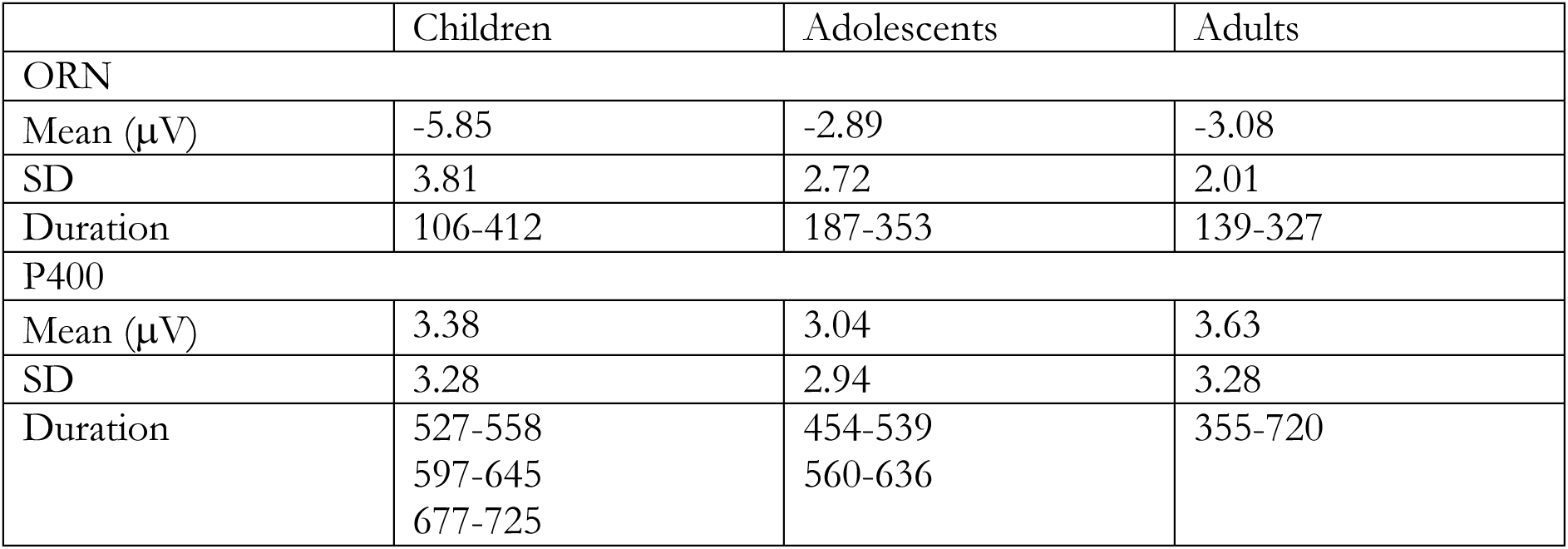
Descriptive statistics of ORN and P400, by age group. Duration represents the periods (>20 ms) of the 70-800 ms post-stimulus epoch (in ms) that contained a significant ORN (70-500 ms) or P400 (300-800 ms) for each age group.

|  | Children | Adolescents | Adults |
| --- | --- | --- | --- |
| ORN |  |  |  |
| Mean (μV) | -5.85 | -2.89 | -3.08 |
| SD | 3.81 | 2.72 | 2.01 |
| Duration | 106-412 | 187-353 | 139-327 |
| P400 |  |  |  |
| Mean (μV) | 3.38 | 3.04 | 3.63 |
| SD | 3.28 | 2.94 | 3.28 |
| Duration | 527-558<br>597-645<br>677-725 | 454-539<br>560-636 | 355-720 |

### General procedure

All participants completed the tasks in the same order. They began with the MICRO-PROMS auditory task, followed by the speech perception task. Then, they participated in the auditory stream segregation task while their electroencephalogram (EEG) was recorded in a soundproof booth. The EEG recording session lasted approximately 90 minutes including setup.

### Statistical Analyses

Statistical analyses were conducted using R Studio software (R Core Team (2021)). First, we assessed the normality of the distribution of behavioral data for each age cohort and dependent variable to guide the choice of statistical models. Data from the streaming task were normally distributed for all groups (Shapiro-Wilk test: *p* = .785 for children, *p* = .628 for adolescents, *p* = .216 for adults). Similarly, data from the SSN condition showed normal distribution for adults (*p* = .454) and children (*p* = .097), while adolescents showed a significant deviation (*p* = .110). However, for the 1-talker condition, data did not meet the assumption of normality for adolescents and adults (*p* < .05), but were normally distributed for children (*p* = .273). No outliers were detected, as all values were within three standard deviations of the mean. We conducted one-sided t-tests to determine whether the performance was above chance/below ceiling in the behavioural tasks. In all three groups of age, participants performed significantly above chance in the streaming task (all *ps* < 0.05), and below ceiling in the speech perception in noise tasks (all *ps* < 0.05).

Next, we assessed developmental effects on behavioural and neurophysiological measures in different ways. Throughout the analyses, we treated age as a continuous variable to better capture developmental changes. In the event of a significant interaction involving age, post-hoc analyses were performed where age was treated categorically.

With respect to the neurophysiological data, we first computed linear regressions to determine whether age, music abilities or their interaction predicted the amplitude of the neurophysiological responses. Then, logistic regressions were performed to evaluate the proportion of participants who presented a significant ORN and P400 as a function of age. Linear regressions were performed again, on the subset of participants who did show a significant ORN/P400 – see supplementary materials. In all models, the factor listener was used as a random intercept.

In addition, we examined the relationship between the neurophysiological data and behavioural auditory performance (streaming and SiN perception). A linear regressions was conducted on stream segregation, with age, musical abilities, amplitude and latency of both ORN, and P400 components, as well as all their interactions (all orders) as predictors. To identify the optimal model and the most relevant interactions, we conducted a LASSO (Least Absolute Shrinkage and Selection Operator) regression. LASSO regression identifies variables and coefficients that minimize prediction error by constraining model parameters. It simplifies high-dimension models by shrinking regression coefficients toward zero, limiting their total sum. Variables with zero coefficients are excluded from the final model. This method reduces model complexity and improves its predictive capability (Ranstam & Cook, 2018). A two-level Generalized Linear Mixed Models (GLMM) was used to explore whether age, musical abilities, stream segregation performance, amplitude of ORN and P400 predicted speech perception in noise (each level of the GLMM represented a noise condition). Here again, LASSO regressions were applied to identify the most relevant interactions and simplify the final model. All statistically significant results are reported with their η² values, an indication of effect size.

## RESULTS

### Development of neural signatures of stream segregation

Figure 1 shows the grand-average of the differential waveforms (figure trials – control trials) evoked in the fronto-central cluster and the centro-parietal cluster in each age group. In all three groups, an early negativity component appears in the fronto-central cluster of electrodes, representing the ORN. It is followed by a positive component with a centro-parietal distribution, reflecting the P400. Table 3 presents the periods of significant ORN/P400 for each age group. At the group level, children displayed earlier and longer ORN than adolescents/adults. The P400 of children and adolescents appears to be shorter and discontinuous compared to adults.

**Table 3:** Results of the linear mixed-effect analyses for the amplitude and latency of ORN and P400.

| Component | Effect | Amplitude |  |  |  | Latency |  |  |  |
| --- | --- | --- | --- | --- | --- | --- | --- | --- | --- |
| | | F | df | p-val | $\eta^2$ | F | df | p-val | $\eta^2$ |
| ORN | **Age** | **8.26** | **1** | **0.005** | **0.11** | **5.27** | **1** | **0.025** | **0.05** |
|  | PROMS | 0.63 | 1 | 0.430 | 0.00 | 2.52 | 1 | 0.117 | 0.03 |
| | Age × PROMS | 0.00 | 1 | 0.966 | 0.00 | 0.89 | 1 | 0.346 | 0.01 |
| P400 | Age | 0.22 | 1 | 0.639 | 0.00 | **12.92** | **1** | **< 0.001** | **0.16** |
|  | PROMS | 0.47 | 1 | 0.496 | 0.00 | 0.01 | 1 | 0.905 | 0.00 |
| | Age × PROMS | 1.29 | 1 | 0.259 | 0.02 | 0.36 | 1 | 0.551 | 0.00 |

Note: The best fitting models for each component amplitude and latency were all an acceptable fit (ORN amplitude [AIC = 360.9, $R^2 = 0.119$ ], latency [AIC = 774.7, $R^2 = 0.116$ ]; P400 amplitude [AIC = 349.7, $R^2 = 0.03$ ], latency [AIC = 833.9, $R^2 = 0.17$ ]). Effects that are significant are shown in boldface.

### ORN

Throughout development, both amplitude and latency of the ORN decreased with age (both *p*s < 0.05); see Table 3 and Figure 2. There was no significant effect of PROMS nor a significant age × PROMS interaction on ORN amplitude or latency (all *p*s > .05). To examine whether the age effect could be due to differences in the variability of the ORN peak latency throughout development, we conducted Levene’s tests for equality of variance. There were no significant differences in variance between the three age groups (*p* > 0.10).

**Figure 2:**
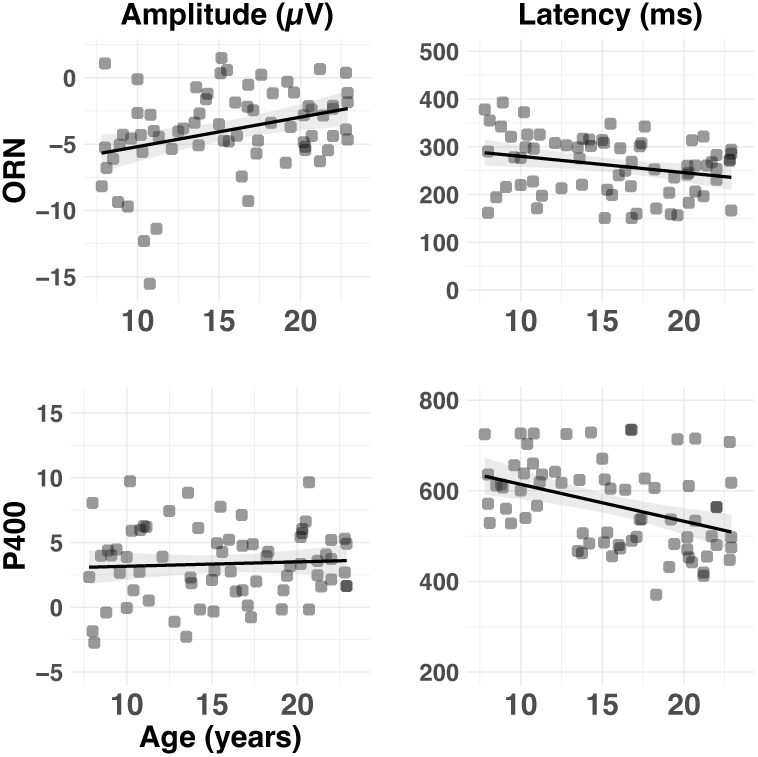
Amplitude (µV) and latency (ms) of the ORN and P400 as a function of age. Individual (dots) and group (lines) data are shown in each panel. Shaded lines represent the 95% confidence interval.

The proportion of participants showing significant ORN at the individual level ranged from 50% (in adolescents) to 72% (in adults) and even 82% (in children), see Figure 3. Despite the surprisingly lower rate observed in adolescents, there was no significant effect of age (treated as a continuous variable) on the participants’ probability to exhibit an ORN (*p* > .10). Linear regressions were performed on the amplitude/latency of the subset of individuals who did show a significant ORN. The same effect of age held significant: ORN amplitude/latency decreased with age (see supplementary Table 1).

**Figure 3:**
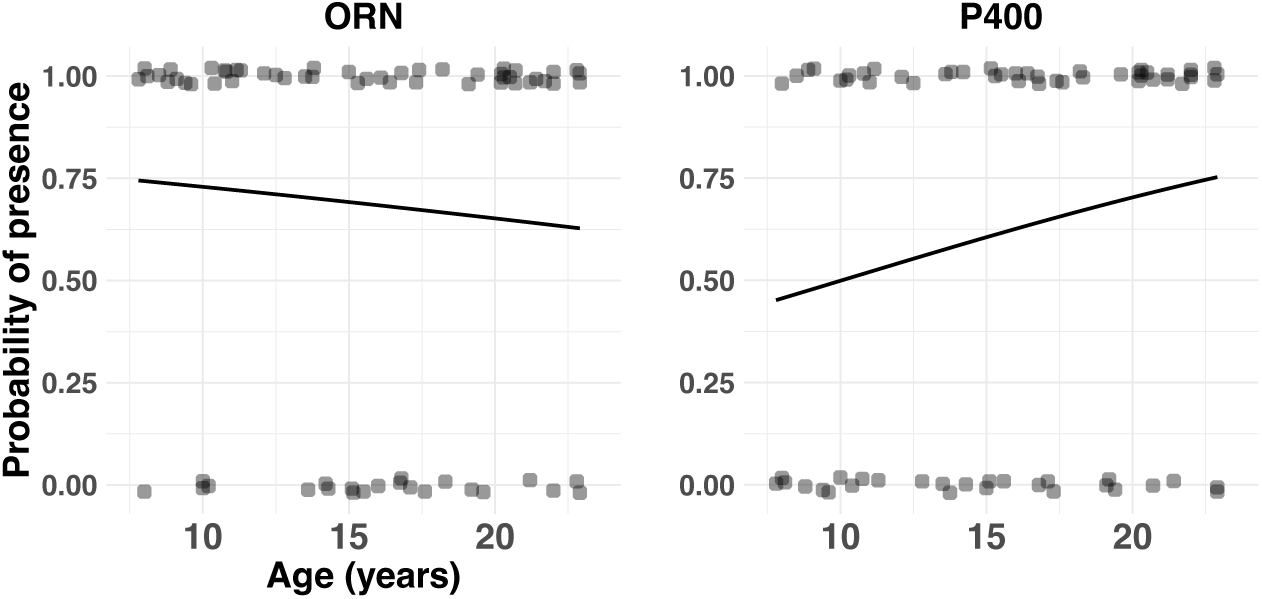
Probability of presence of ORN and P400 as a function of age. Individual (circles) and group (lines) data are shown for both components. At the individual level, components were either present (probability = 1) or absent (probability = 0). Solid lines results from a logistic regression fitting a general linear model with age as a predictor and presence of the component as the outcome variable.

### P400

There was no significant effect of age, PROMS nor age × PROMS interaction on P400 amplitude (all *p*s > .10); see Table 3 and Figure 2. P400 latency significantly decreased with age (*p* < 0.001). Variability in P400 latency was larger in children than in adolescents and adults, as indicated by Levene’s test for equality of variance (both *p*s < 0.05). There was no significant difference in variance of P400 latency between adolescents and adults (*p* > 0.10).

The proportion of participants showing significant P400 at the individual level ranged from 52% (in children) and 72% (in adults), see Figure 3. There was no significant effect of age (treated as a continuous variable) on the participants’ probability to exhibit a P400 (*p* > .10). The same age effects held significant when analyses were conducted only on the subset of individuals who showed a significant P400 (see supplementary Table 1).

### Predictors of stream segregation performance

What predictors contribute to the performance in figure-ground segregation? To answer this question, we performed a least absolute shrinkage and selection operator (LASSO) regularized regression, which enforces sparse solutions. The initial selection indicated the age × PROMS × ORN amplitude × P400 amplitude as a potential interaction to investigate. A subsequent linear regression analysis was thus conducted with these four factors and all their 2-, 3- and 4-way interactions as predictors of the figure-ground performance. Results reveal two significant main effects, each illustrated in Figure 4 (also see Supplementary Table 2). The main effect of age indicates that performance improves from childhood to adulthood (*p* < 0.001). Notably, P400 (but not ORN) amplitude also predicted figure-ground segregation: larger P400 responses led to higher dʹ (*p* < 0.001). There was also a marginally significant effect of PROMS: better musical abilities were associated to higher dʹ on the segregation task (*p* = 0.053). Note that there was no significant effect of age on PROMS (*p* > .10).

**Figure 4:**
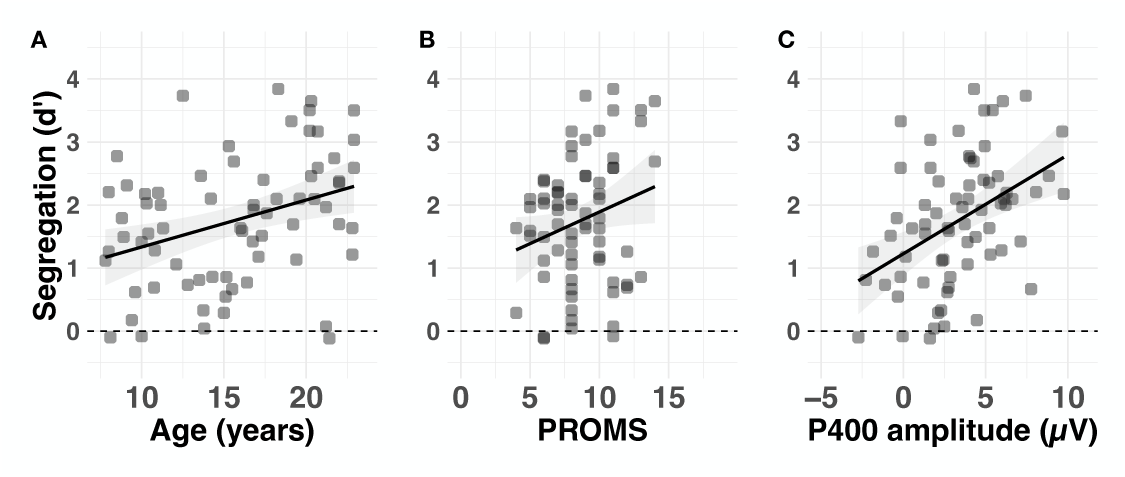
Predictors of the behavioural stream segregation (dʹ) performance. **A:** Age (in years), **B:** PROMS and **C:** P400 amplitude (in µV). Dashed horizontal line indicates chance level.

### Predictors of SiN performance

LASSO regularized regression was also applied to identify the potential factors contributing to speech perception in noise performance. It indicated that the triple interaction noise condition × ORN latency × P400 latency as well as the age × stream segregation performance could be predictors of the SiN perception. A subsequent generalized logistic regression was conducted with these two interactions (as well as the main effects associated with each interaction) as predictors of SiN perception (for details, see Supplementary Table 3). Significant findings are illustrated in Figure 5. The main effect of condition was significant: overall, listeners performed better in the presence of one interfering speaker than in SSN (*p* = 0.004). With respect to the interactions, only the age × stream segregation came out significant (*p* = 0.008). Post-hoc analyses indicate that in adults, better stream segregation predicted better SiN perception (*p* < 0.001). This was not the case in either children or adolescents (both *ps* > 0.10).

**Figure 5:**
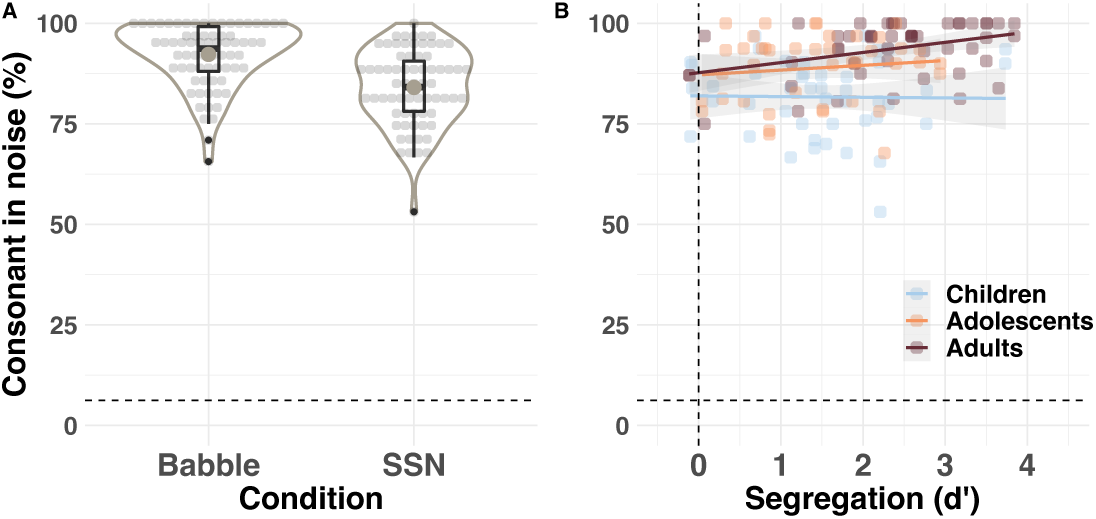
Predictors of the consonant identification in noise. **A**: noise condition, B: Interaction between stream segregation (dʹ) and age. The interaction was decomposed and is illustrated by age band. The dashed lines indicated chance level.

## DISCUSSION

Neurophysiological responses (ORN and P400) were recorded in children, adolescents and young adults, as they performed a stream segregation task based on temporal coherence. Both amplitude and latency of the ORN decreased between childhood and early adulthood. P400 amplitude did not significantly change, yet P400 latency also decreased throughout development. Notably, P400 amplitude significantly predicted behavioural performance in the stream segregation task (together with age). In turn, stream segregation significantly predicted consonant identification in noise in adults, but not in children or adolescents.

### Neurophysiological measures

At the group level, children, adolescents and adults all showed a significant ORN, followed by a significant (albeit ‘segmented’) P400. This contrasts with previous studies conducted in school-age children, showing a significant ORN but no P400 (Alain et al., 2003; Mehrkian et al., 2022). This discrepancy is likely attributable to differences in the electrode cluster selected to analyse the P400: previous children studies focused on a cluster of fronto-central electrodes, whereas we analysed centro-parietal electrodes where the P400 is typically maximal in adults (Tóth et al., 2016). This finding indicates that children, like adolescents and adults, do exhibit the neural markers of both bottom-up (ORN) and top-down (P400) processing of stream segregation based on temporal coherence.

Both ORN and P400 appear to follow protracted developmental trajectories, with peak latencies gradually decreasing from childhood to adulthood. This developmental latency decrease is small-to-medium for the ORN, and strong for the P400. Many developmental ERP studies report decreasing latencies throughout the childhood and adolescence years (Oades et al., 1997; Gomot et al., 2000; Bishop, Hardiman and Barry, 2011). Shorter latencies are generally interpreted as indicative of faster processing speed, which has been related to the myelination of the neural tracts linking generators of the different ERP components (Huttenlocher, 1979; for a review see Musiek, Verkest and Gollegly, 1988). Axon myelination supports maturation of the human connectome by means of increased speed and fidelity of neural signalling. In humans, myelination follows an exceptionally long developmental trajectory, extending beyond late adolescence (Corrigan et al., 2021; Miller, Stimpson & Sherwood, 2012; Rowley et al., 2017). This extended maturation of the white matter in thalamo-cortical tracts likely explains the decrease in ORN and P400 latency throughout adolescence.

In parallel to shorter latencies, ORN amplitude gradually decreased from childhood to adulthood – in line with previous results collected in children (Alain et al., 2003). Two (non-mutually exclusive) interpretations are possible. Cortical ERPs reflect the aggregate measure of post-synaptic potentials of pyramidal cells within a given population of neurons in response to a stimulus. In adults, both ORN and P400 are thought to reflect activity of generators located in the primary auditory and anterior cingulate cortices (Bharadwaj et al., 2022; Tóth et al., 2016). Additionally, the ORN has specific generators in the superior temporal gyrus and in the inferior parietal sulcus (Molloy, Lavie and Chait, 2019; Tóth et al., 2016). ORN generators thus appear mostly localised in the temporal and frontal areas of the brain. Coincidentally, synaptic pruning accelerates around puberty onset in frontal, temporal and parietal brain areas (Vijayakumar et al., 2021a, Vijayakumar et al., 2021b). In fact, synaptic pruning is thought to relate to a decrease in the spectral composition of resting EEG below 8 Hz during the adolescent period (Whitford et al., 2007). A first explanation to the decreasing ORN amplitude throughout development might thus pertain to synaptic pruning at adolescence.

Second, age differences in ORN amplitude might be attributable to structural changes to the generators of the response. The lower probability of adolescents to show a significant ORN at the individual level could support this interpretation (even though it did not reach significance). The EEG is thought to best pick activity from neurons that are oriented perpendicular to the cortical surface, forming current dipoles (Nunez & Srinivasan, 2006). Structural changes could thus impact the orientation of the current dipoles, affecting the adolescent and adult surface ERPs, compared to the children’s. We did not observe obvious changes in the distribution of amplitude of the ERP across the scalp throughout development. Studies using magnetoencephalography are warranted to further our understanding of the developmental changes in generators of the ORN, and their relationship to ORN amplitude.

Such studies should also seek to explain why we observed a developmental decrease in the ORN, but not in the P400 amplitude. Noteworthily, children’s P400 peak latency was significantly more variable than adolescents’ and adults’. This could explain the smaller (and more ‘segmented’) P400 response observed in our youngest participants. What factors drive the larger variability in P400 peak latency during childhood remains an open question, certainly worth investigating in the future.

### Behavioural measures

We identified two significant main predictors of stream segregation based on temporal coherence: age and P400 amplitude recorded during the segregation task. The medium effect of age indicates a gradual improvement of stream segregation, from childhood to adulthood. So far, behavioural results indicated persistent immaturity in auditory streaming until at least 12 years of age, in paradigms that rely on either sequential (Sussman et al., 2007; Sussman & Steinschneider, 2009; Sussman et al., 2015; Sussman et al., 2001; Lepisto et al., 2009) or concurrent stream segregation (Alain et al., 2003; for a review, see Calcus, 2024). Our results confirm this protracted development of stream segregation until adulthood in complex auditory scenes where streaming must operate based on the temporal coherence across auditory filters (Shamma, Elhilali and Micheyl, 2011).

Irrespective of the listeners’ age, perceptual musical abilities measured in non-musicians tend to predict stream segregation. In adults, musical expertise has been linked to improved stream segregation (Alain et al., 2014; Zendel et al., 2009; Zendel & Alain, 2014). Our results suggest that these observations might not be related to musical training per se, but to inherent musical abilities. Longitudinal studies are needed to better understand the transfer effect of musical expertise to auditory abilities.

Last but not least, P400 amplitude strongly predicts performance on the streaming task in our sample of participants (irrespective of age). This concurs with the existing adult literature (Alain et al., 2001; Tóth et al., 2016), and extends the findings to children and adolescents. In other words, even though we did not find a significant age effect on the P400 amplitude, this component appears to predict behavioural performance (which itself does improve with age). The P400 thus appears to reflect attentional and decisional processes involved in stream segregation, already in childhood.

Interestingly, stream segregation performance was shown to predict SiN (whatever the condition), but only in adults: this relationship was not significant in either children or adolescents. This supports previous reports of a marginally significant relationship between figure detection in a stochastic background and speech-in-babble in adults (Holmes & Griffiths, 2019). In our recent study (Benocci & Calcus, 2024), we used structural equation modelling to investigate the relationship between age (8-22 years), musical abilities, stream segregation and SiN perception – all tested with material that was very similar to that used in the present study. The results indicated a significant effect of age on stream segregation, which itself contributed to SiN perception. In this paper, we did not report the results of the linear regression conducted on SiN perception with age and stream segregation as predictors. In the spirit of replication, we have now re-analysed this earlier dataset, and observe a significant relationship between stream segregation and SiN perception in adults (p = .026). This relationship was marginally significant in adolescents (p = .075), and was not significant in children (p = .667). Therefore, the relationship between stream segregation and SiN might only appear gradually throughout adolescence. Children and adolescents may not yet be able to rely on their immature stream segregation abilities to process SiN efficiently. Future work should seek to determine whether children/adolescents rely on other (auditory, linguistic or cognitive) factors to perform SiN, and/or whether the relationship between streaming and SiN emerges as a result of exposure to complex auditory backgrounds.

To sum up, SiN perception is predicted by the behavioural streaming performance in adults – but not in children or adolescents. Stream segregation itself improves gradually from childhood to adulthood, and is better in individuals who show larger P400 (irrespective of their age). Unusual as it might be, we would like to highlight the factors that do *not* appear to contribute to SiN perception in our study: none of the neurophysiological markers of stream segregation turned out to be significant SiN predictors. This observation appears to contradict a number of studies that found significant relationships between (cortical or subcortical) ERPs and SiN perception throughout the lifespan (Anderson et al., 2010a; Anderson et al., 2010b; Anderson, et al., 2010c; Anderson et al., 2011; Anderson et al., 2013; Koerner et al., 2016; Parbery-Clark et al., 2011; Thompson et al., 2019). Our data suggest that, in adults, the only significant predictor of SiN perception is stream segregation, as measured by a task that is closely related to SiN. Performance in basic auditory scene analysis may thus present a bottleneck to SiN difficulties in children and adolescents.

### Clinical relevance

Children have poorer SiN perception than adults (for a review, see Leibold, 2019). Adding insult to injury, classrooms are often very noisy. Whereas this double penalty is distressing for all children, some of them are disproportionately affected by the presence of background noise. In 2021, close to 100 million children and adolescents (< 20 years) were affected by hearing loss worldwide (Guo et al., 2024). Even the milder degrees of hearing loss affect SiN perception (Crandell, 1993; Lewis, Valente & Spalding, 2015; Goldsworthy & Markle, 2019). Childhood hearing loss is known to affect the central processing of sounds, with some effects that only reveal themselves at adolescence (Calcus et al., 2019). Children with a hearing loss also show smaller/later ORN than age-matched children with normal hearing (Mehrkian et al., 2022). How a peripheral hearing loss affects the maturation of P400, stream segregation performance and its late-emerging relationship with SiN remains an open question.

Some children report having listening difficulties despite normal hearing (as measured by the audiogram), a condition sometimes referred to as ‘auditory processing disorder’ (APD, for a review, see Moore & Hunter, 2013). Children with APD experience pronounced difficulties with SiN and often exhibit impaired stream segregation abilities (Cameron & Dillon, 2008; Lagacé, Jutras and Gagné, 2010; Moosavi et al., 2015). Whether their difficulties involve the bottom-up, automatic processing of streaming (ORN), and/or the top-down, attentional processing of the streams (P400) – and how these two components develop in the context of APD - has remained so far unexplored.

In general, children with developmental disorders (e.g., ADHD, reading difficulties) also perform more poorly in SiN tasks than typically developing children (Blomberg et al., 2019; Calcus, et al., 2018; Ziegler et al., 2005; Ziegler et al., 2009). Better characterizing SiN difficulties throughout development, and their relationship to stream segregation, could open new avenues for diagnosis and intervention.

### Limitations

Our SiN task was designed to be quick and easy to administer to young participants, and to somewhat resemble the streaming task (similar duration, similar auditory signal to detect amidst the background noise). Therefore, we used a consonant identification task in noise, at a fixed SNR, in the presence of only one interfering talking and in SSN. This led to a very good (non-normally distributed) performance in the 1-talker condition, even though it was significantly lower than ceiling. It may be the case that the relationship between the streaming task and SiN is different with different types of material and measure. Future studies would benefit from measuring speech reception thresholds and incorporating more ecological stimuli that vary in e.g. linguistic complexity, sound source spatialization, and number of interfering talkers.

Additionally, ERPs were only recorded in an active condition, as participants performed the stream segregation task. Including a passive listening condition could clarify the role of automatic versus controlled auditory processing on stream segregation. For example, children and adults are thought to have similar MMN amplitude if they are actively engaged in the streaming task, but not if they listen passively to the sequences (Sussman & Steinschneider, 2009).

## Conclusion

In conclusion, our results reveal a progressive maturation of both automatic and attention-driven mechanisms underlying stream segregation, extending until adulthood. P400 amplitude emerges as a strong predictor of segregation performance, highlighting the critical role of attentional processes on auditory scene analysis across development. Notably, the relationship between stream segregation and SiN perception only manifests in adulthood, suggesting a late maturation of the integration between these processes. Basic mechanisms of non-linguistic stream segregation may thus present a bottleneck to SiN difficulties in children and adolescents. Therefore, our findings carry significant clinical implications, in particular for children with peripheral and/or central impairments along the auditory pathway.

## ACKNOWLEDGMENTS

We are very grateful for the time and efforts of the children who participated in this study and to their parents. We would like to thank Luna Leonardy, Hasnae El Arround, Jessy Angama-Gravier, Mélanie Rupaire, Aubrey Plume and Manon Couvignou for their assistance with recruitment and data collection. This work was supported by a grant from The Belgian Kids’ Fund for Pediatric Research awarded to Elena Benocci, and by a European Research Council Starting Grant “SensationaHL” (101076968) awarded to Axelle Calcus.

