## SupplementaryMaterial for "Neural signatures of stream segregation: From childhood to adulthood"

**Supplementary material**

Supplementary Table 1: Results of the linear regressions conducted on the amplitude and latency of ORN and P400, where present.

| | | Amplitude | | | | Latency | | | |
|---|---|---|---|---|---|---|---|---|---|
| Component | Effect | F | df | p-val | $\eta^2$ | F | df | p-val | $\eta^2$ |
| ORN | **Age** | **10.38** | **1** | **0.002** | **0.20** | **6.28** | **1** | **0.016** | **0.07** |
| | PROMS | 0.63 | 1 | 0.430 | 0.00 | **6.39** | **1** | **0.015** | **0.11** |
| | Age × PROMS | 0.00 | 1 | 0.966 | 0.00 | 1.60 | 1 | 0.212 | 0.02 |
| P400 | Age | 2.82 | 1 | 0.101 | 0.07 | **8.61** | **1** | **0.005** | **0.17** |
| | PROMS | 0.13 | 1 | 0.714 | 0.00 | 0.08 | 1 | 0.770 | 0.00 |
| | Age × PROMS | 1.14 | 1 | 0.711 | 0.00 | 1.62 | 1 | 0.209 | 0.03 |

Note: The best fitting models for each component amplitude and latency were all an acceptable fit (ORN amplitude [AIC = 230.4, $R^2$ = 0.213], latency [AIC = 512.1, $R^2$ = 0.245]; P400 amplitude [AIC = 189.4, $R^2$= 0.07], latency [AIC = 506.9, $R^2$= 0.21]). Effects that are significant are shown in boldface.

Supplementary Table 2: Results of the linear regression conducted on discrimination performance (d′) in the stream segregation task

| | F | df | p-val | $\eta^2$ |
|---|---|---|---|---|
| **Age** | **12.53** | **1** | **<0.001** | **0.08** |
| PROMS | 3.88 | 1 | 0.053 | 0.05 |
| **P400 amplitude** | **19.1** | **1** | **<0.001** | **0.16** |
| Age × PROMS | 3.35 | 1 | 0.072 | 0.03 |
| Age × P400 amp | 0.05 | 1 | 0.823 | 0.00 |
| PROMS × P400 amp | 0.39 | 1 | 0.534 | 0.00 |
| Age × PROMS × P400 amp | 0.73 | 1 | 0.393 | 0.00 |

Note: The model was an acceptable fit [AIC = 182.5, $R^2$ = 0.392]. Significant effects are shown in boldface.

Supplementary Table 3: Results of the logistic regression conducted on speech perception in noise scores (%)

| | coefficient | std error | df | p-val | $\eta^2$ |
|---|---|---|---|---|---|
| **Condition** | **-1.74** | **0.61** | **1** | **0.004** | **0.19** |
| P400 latency | -0.00 | 0.00 | 1 | 0.119 | 0.00 |
| Age | 0.02 | 0.03 | 1 | 0.352 | 0.71 |
| d′ segregation | -0.43 | 0.23 | 1 | 0.064 | 0.17 |
| Condition × P400 latency | 0.00 | 0.00 | 1 | 0.145 | 0.00 |
| **Age × d′ segregation** | **0.04** | **0.01** | **1** | **0.009** | **0.00** |

Note: The model was an acceptable fit [AIC = 557.5]. Significant effects are shown in boldface.
